# *In silico* and *in vitro* screening of FDA-approved drugs for potential repurposing against tuberculosis

**DOI:** 10.1101/228171

**Authors:** Sridharan Brindha, Jagadish Chandrabose Sundaramurthi, Savariar Vincent, Devadasan Velmurugan, John Joel Gnanadoss

**Affiliations:** Loyola College, Nungambakkam, Chennai 600034, Tamil Nadu, India; National Institute for Research in Tuberculosis (ICMR), Chetpet, Chennai 600031, Tamil Nadu, India; CAS in Crystallography and Biophysics, University of Madras, Guindy Campus, Chennai – 600025, Tamil Nadu, India

**Keywords:** Tuberculosis, Repurposing, *trpD*, coaA, Lymecycline, Cefpodoxime

## Abstract

**Motivation:** Repurposing of known drugs to newer clinical conditions is a promising avenue for finding novel therapeutic applications for tuberculosis.

**Methods:** We performed docking-based virtual screening for 1554 known drugs against two of the potential drug targets, namely *trpD* and coaA of *M. tuberculosis.* In the first round of *in silico* screening we used rigid docking using Glide and AutoDock Vina. We subjected the consistently ranked drugs for induced-fit docking by these tools against the same target proteins. We performed luciferase reporter phage (LRP) assay to determine the biological activity of five selected drugs against *M. tuberculosis.*

**Results:** We observed lymecycline and cefpodoxime to be active against drug susceptible and drug resistant strains of *M. tuberculosis.* In addition, lymecycline and cefpodoxime showed synergistic activity with rifampin and isoniazid against *M. tuberculosis.*

**Conclusion:** Our results suggest that lymecycline and cefpodoxime have potential to be repurposed for the treatment of tuberculosis.

## 1 INTRODUCTION

Tuberculosis is the world leading infectious killer disease accounting for 1.3 million deaths and an additional 374 000 deaths among HIV-positive people globally in 2016 (WHO 2017). Though tuberculosis can be cured using the currently available drugs, emergence of drug resistant *Mycobacterium* tuberculosis that results in multiple forms drug resistant TB including multidrug-resistant tuberculosis (MDR-TB) and extensively drug-resistant tuberculosis (XDR-TB) necessitates the discovery of newer therapeutics for the sustained success in the treatment and control of TB. Another difficulty in the management of TB is that minimum of six months is required for the treatment with rifampin, isoniazid, ethambutol and pyrazinamide for first four months and rifampin and isoniazid for next two months. The present situation of the TB treatment certainly demands for much more effective drugs with which we can treat the TB in significantly shorter time and lesser doses.

Conventional process of drug discovery needs about 10 years or more and a big budget of up to billion dollars. In this background, identification of newer therapeutic uses for already known drugs against other diseases, which is referred as repurposing is considered to be a promising avenue in the pharmaceutical industry and academia (Ashburn and Thor 2004). Repurposing is also known by several other terminologies viz., redirecting, repositioning, or reprofiling while identifying new clinical use for molecules which failed during the clinical development stage for its original goal is referred as drug rescue (Langedijk et al., 2015). The scientific principle of repurposing of drugs is that some of the drugs can interact with multiple drug targets. Polypharmacology is a term used to explain the ability of those drugs which can act simultaneously and specifically on multiple targets (Anighoro et al., 2014). Drugs with polypharmacological nature have relatively lower chances for the development of drug resistance since they target multiple drug targets, and show better efficacy compared to drugs with specificity for a single target, especially against advanced-stage diseases (Anighoro et al., 2014; Peters, 2013). However, the polypharmacology differs from the unintended promiscuousity of drugs with unknown target that leads to adverse reactions (Anighoro et al., 2014).

Several drugs have already been successfully repurposed for new diseases, outside the original medical scope. For example, metformin is a commonly used drug for the treatment of type 2 diabetes, now repurposed for cancer (Kasznicki et al., 2014), colesevelam, used for hyperlipidemia and repurposed for type-2-diabetes (Finsterer and Frank, 2013), Gabapentin used for epilepsy, now repurposed for neuropathic pain (https://www.drugbank.ca/drugs/DB00996). Drugs which failed to meet the intended purpose can also be repurposed for the treatment of other diseases; for example, zidovudine was first synthesized as a potential anti-cancer agent in 1964 but failed in the cancer treatment, and later in 1984 it was discovered to be active against HIV, and subsequently approved for the treatment in 1987 (Broder, 2010). Recognizing the potential of repurposing of drugs in discovery of newer therapeutic applications especially against diseases that has no drugs to treat or difficult treat, recently government agencies viz., National Institute of Health (USA) and Medical Research Council (UK) established collaborative research with private pharmaceutical industries to promote the repurposing of drugs (Frail et al., 2015).

Several drugs, including, fluroquinolones, clofazimine, meropenem and clavulanate were primarily used for other clinical conditions and now repurposed successfully for treatment of drug resistant forms of TB (Palomino and Martin 2013; Maitra et al., 2015). In addition, several more drugs, to name a few, verapamil (Adams et al., 2011, Gupta et al., 2013 and Adams et al., 2014), statins, (Skerry et al., 2014, Parihar et al., 2014 and Lobato et al., 2014) and metformin (Singhal et al., 2014) have been reported to have potential use in the treatment of TB. These evidences suggest that repurposing of old drugs could be a promising avenue for finding a newer therapeutic drugs for TB.

The genome of *M. tuberculosis* encodes for about 4000 proteins which necessitates scrupulous selection of drug targets for drug discovery. Anthranilate phosphoribosyltransferase (*trpD*, Rv2192c) is known to be essential for the optimal growth of *M. tuberculosis* (Sassetti et al., 2003); it was also demonstrated that the *M. tuberculosis* auxotrophic for tryptophan (Smith et al., 2001) and the *trpD* is involved in the biosynthesis of phenylalanine, tyrosine and tryptophan in *M. tuberculosis* (http://www.genome.jp/kegg/). Pantothenate kinase (*coaA/panK* Rv1092c) catalyzes the first step of the universal coenzyme A biosynthetic pathway in *M. tuberculosis* (Das et al., 2006; Ref:http://www.uniprot.org/uniprot/P9WPA7) and it is an essential gene for the optimal growth of *M. tuberculosis* (Sassetti et al., 2003; Awasthy et al., 2010) and survival during infection (Sassetti and Rubin, 2003). Further, several bioinformatics analysis revealed *trpD* (Defelipe et al., 2016; Kushwaha and Shakya 2010; Anishetty et al., 2005) and coaA (Raman et al., 2008; Defelipe et al., 2016) as potential drug targets for *M. tuberculosis.* Both *trpD* and *coaA* of *M. tuberculosis* lack homolog proteins in the human genome. For both of these two proteins, three dimensional structures are available in the Protein Data Bank (Castell et al., 2013 and Chetnani et al., 2010). All these factors encouraged us to select *trpD* and *coaA* as target proteins against which molecular docking was performed.

In the present study we undertook an effort to prioritize 1554 FDA approved drugs using docking-based virtual screening against two of the potential drug targets and tested some of the prioritized drugs using Luciferase Reporter Phage (LRP) assay against *M. tuberculosis*. We identified cefpodoxime and lymecycline to inhibit drug susceptible and resistant strains of *M. tuberculosis*, and have synergistic role with two of the major anti-tubercular drugs, rifampin and isoniazid.

## 2 METHODS

### 2.1 *In silico* prioritization of drugs

A total of 1554 FDA-approved drugs were obtained from DrugBank and virtually screened against two proteins namely anthranilate phosphoribosyltransferase (*trpD* - Rv2192c) and pantothenate kinase (*coaA* - Rv1092c) of *M. tuberculosis* using molecular docking with the help of two different docking tools, namely Glide and AutoDock Vina. Rigid body docking algorithm was employed for initial screening. Those drugs which were ranked in top 10% during rigid-body docking by both Glide and ADV were subjected for induced-fit docking algorithm for further prioritization. Figure 1 and 4 display the workflow of the procedure.

**Figure 1:**
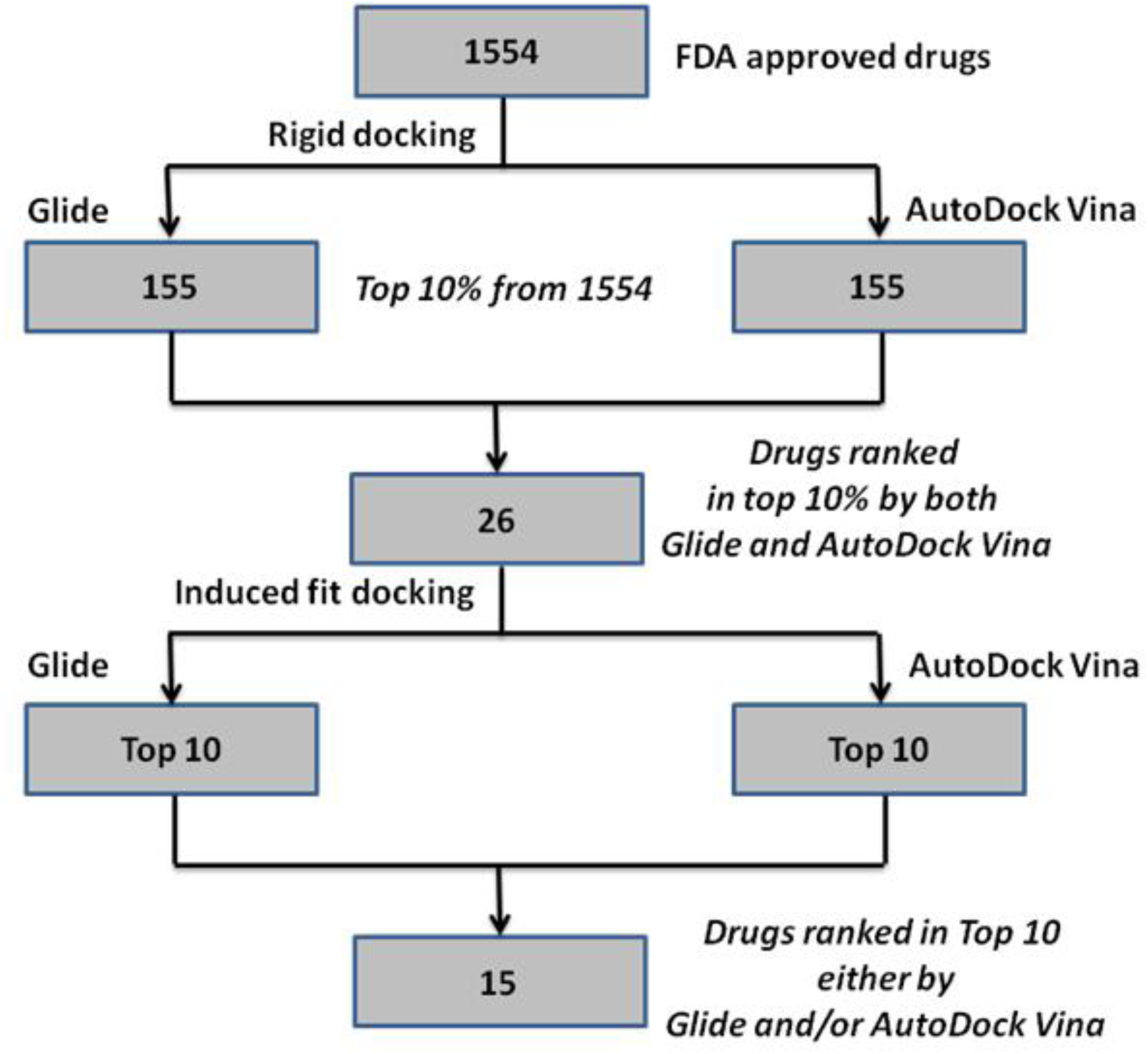
Prioritization of 1554 known drugs using virtual screening against *trpD* of *M. tuberculosis*.

### 2.2 In vitro *testing by LRP assay against* M. tuberculosis

Biological activity of five of the prioritized drugs, namely, cefpodoxime, montelukast, lymecycline, diosmin and carboprost tromethamine were evaluated against *M. tuberculosis* using LRP assay (Kumar et al., 2008; Dusthackeer et al., 2008; Manikkam et al., 2014). Initially, all of the drugs were tested against drug sensitive *M. tuberculosis* H37Rv strain at two different concentrations (20 μg/ml and 100μg/ml). Those drugs which were active in the initial screening were further tested against multidrug-resistant strains of *M. tuberculosis* at four different concentrations (10, 20, 50 and 100μg/ml). Additionally, the active drugs were also tested for synergistic effect with two of the major first-line anti-tubercular drugs rifampin and isoniazid using LRP assay.

## 3 RESULTS

### 3.1 Prioritized drugs against *trpD*

We found a total of 26 drugs to be ranked consistently at the end of rigid docking of 1554 FDA approved drugs against *trpD*. When we subjected these 26 drugs for induced-fit docking against *trpD*, 15 drugs were observed to be ranked within top 10 ranks by either Glide or AutoDock Vina (Table 1). Experimentally studied compound 2,2’-iminodibenzoic acid in complex with *trpD* (PDB ID: 3QQS) was used as the control molecule and docked along with *trpD*. All of the 15 prioritized drugs were found to have better binding affinity score by ADV than the control molecule though only top three ranked molecules were observed to have better Glide score than the control molecule (Table 1). Among these 15 shortlisted drugs, five drugs, namely, acarbose (Glide rank 1 and ADV tank: 2), cefpodoxime (2, 5), montelukast (6, 6), lymecycline (7, 1) and idarubicin (10, 7) were ranked consistently within the top 10 drugs by both Glide and ADV.

**Table 1.**
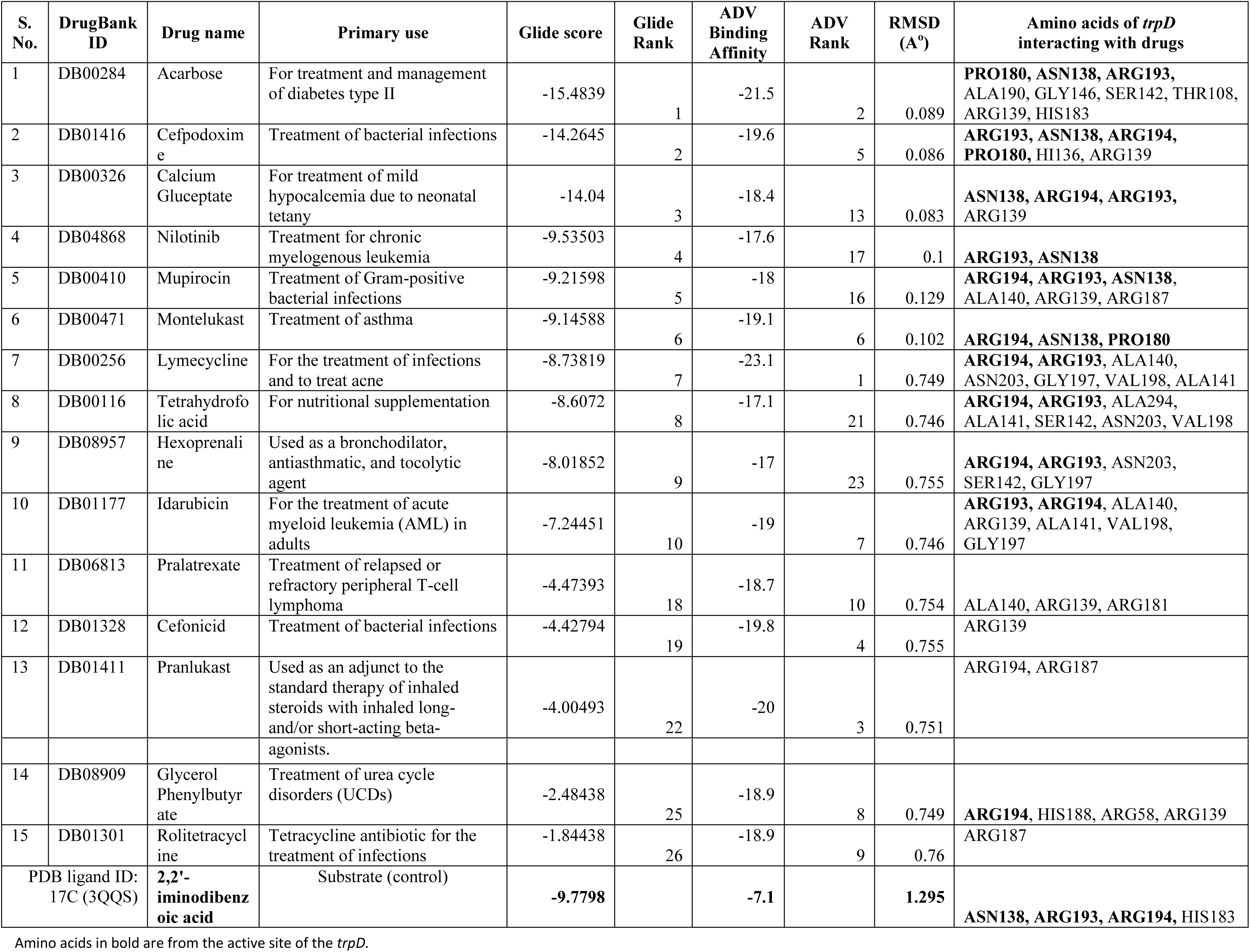
Prioritized drugs which show strong binding interactions with *trpD* of *M. tuberculosis*

### 3.2 Prioritized drugs against *coaA*

When 1554 drugs were screened against *coaA* of *M. tuberculosis*, we observed 16 drugs to be ranked consistently within top 10% in the rigid docking by both Glide and ADV. When we subjected these 16 drugs for induced fit docking by both Glide and ADV, 14 drugs were observed within top 10 ranks (Table 2). Phosphopantothenate (PAZ (N-[(2R)-2-hydroxy-3,3-dimethyl-4-(phosphonooxy)butanoyl]-beta-alanine), one of the ligand resolved in complex with coaA (PDB ID: 3AEZ) was used as the control molecule during docking. All of the 14 drugs were observed to have better binding affinity score with *coaA* than the control molecule, PAZ by ADV while only 8 drugs scored better than PAZ by Glide (Table 2). A total of six drugs namely, flavin adenine dinucleotide (Glide rank: 3, ADV rank: 3), carboprost tromethamine (4, 6), montelukast (5, 9), paromomycin (7, 5), calcium gluceptate (8, 10), lymecycline (9, 4) were found to be consistently ranked within top 10 drugs by both Glide and ADV.

**Table 2.** Prioritized drugs which show strong binding interactions with *coaA* of *M. tuberculosis*

| S. No | Drug Bank ID | Drug name | Primary use | Glide score | Glide Rank | ADV Binding Affinity | ADV Rank | RMSD (Å) | Amino acids of *coaA* interacting with drugs |
| --- | --- | --- | --- | --- | --- | --- | --- | --- | --- |
| 1 | DB08995 | Diosmin | Treatment of venous disease | -13.5027 | 1 | -10.9 | 16 | 0.906 | **HIS179, TYR182, THR128, TYR153**, LYS103, SER104, ALA100, ARG238 |
| 2 | DB00694 | Daunorubicin | For remission induction in acute nonlymphocytic leukemia | -11.9108 | 2 | -11.6 | 14 | 0.898 | **HIS179**, PHE254, ARG108, SER104, LEU40 |
| 3 | DB03147 | Flavin adenine dinucleotide | Ophthalmic treatment for vitamin B2 deficiency | -11.9008 | 3 | -16.8 | 3 | 0.901 | **HIS179**, ALA100, ARG38, LEU40, SER104, THR105, ARG238, ASP129, LYS103, VAL101, GLY102 |
| 4 | DB00429 | Carboprost Tromethamine | A nonsteroidal abortifacient | -11.2632 | 4 | -12.7 | 6 | 0.896 | ASP129, SER104, THR105, ARG108, LYS103, GLY102 |
| 5 | DB00471 | Montelukast | Treatment of asthma | -11.2397 | 5 | -12 | 9 | 0.909 | **TYR182**, PHE254, ARG238, GLY102, LYS103 |
| 6 | DB00691 | Moexipril | Treat high blood pressure | -11.131 | 6 | -11.8 | 12 | 0.9 | **HIS179**, ARG108, GLY41, SER104, THR105, LYS103, GLU42 |
| 7 | DB01421 | Paromomycin | Treatment of acute and chronic intestinal amebiasis | -11.1175 | 7 | -12.9 | 5 | 0.908 | **HIS179**, GLU201, ASP129, SER104, GLY36 |
| 8 | DB00326 | Calcium Gluceptate | Treatment of mild hypocalcemia | -9.98464 | 8 | -12 | 10 | 0.902 | **HIS179**, ARG238, LYS103, ALA100, LEU40, SER104, GLY102 |
| 9 | DB00256 | Lymecycline | For the treatment of infections and to treat acne | -9.65968 | 9 | -13 | 4 | 0.904 | **HIS179**, ARG108, ASP129 |
| 10 | DB01328 | Cefonicid | Treatment of bacterial infections | -9.31335 | 10 | -11.8 | 13 | 0.898 | **ARG108**, LEU40, SER104, LYS103, GLY102, ASP129 |
| 11 | DB01232 | Saquinavir | HIV protease inhibitor | -8.49036 | 12 | -17.5 | 2 | 0.911 | **HIS179, TYR153, TYR182, LEU203**, TYR235, ARG238, LYS103, ASN277 |
| 12 | DB00212 | Remikiren | For the treatment of hypertension and heart failure | -8.45254 | 13 | -12.6 | 7 | 0.906 | **HIS179**, PHE254, ARG238, THR127, GLU201, ARG108 |
| 13 | DB06590 | Ceftaroline fosamil | Treatment of bacterial infections | -8.06396 | 14 | -19.7 | 1 | 0.904 | TYR235, ASN277, LYS103, ARG238 |
| 14 | DB00950 | Fexofenadine | Treatment of hayfever | -7.32914 | 15 | -12.4 | 8 | 0.897 | SER104, THR105, ASP129, LEU40 |
| PAZ |  | Phosphopantothenate |  | **-9.76722** |  | **-5.8** |  | **1.52** | **TYR182, TYR153**, SER104, LYS103, TYR235, ASN277 |

Amino acids in bold are from the active site of the *coaA*.

### 3.3 Lymecycline and cefpodoxime inhibit both drug susceptible and resistant strains of *M. tuberculosis*

Initial *in vitro* screening for activity was carried out at two different concentrations (10, 20, 50 and 100 μg/ml) against drug sensitive strains of *M. tuberculosis*. We found two drugs namely cefpodoxime and lymecycline to inhibit *M. tuberculosis* H37Rv. Cefpodoxime inhibited drug sensitive strains of *M. tuberculosis* at 20, 50 and 100 μg/ml concentrations and lymecycline at 10, 20, 50 and 100 μg/ml concentrations (Table 3). Anti-mycobacterial activity against MDR stains of *M. tuberculosis* was carried out at 20 and 100 μg/ml. Both cefpodoxime and lymecycline inhibited MDR strains at 20 and 100 μg/ml concentrations. Other three drugs tested namely, montelukast, diosmin and carboprost tromethamine did not show inhibitory activity against *M. tuberculosis.*

**Table 3:**
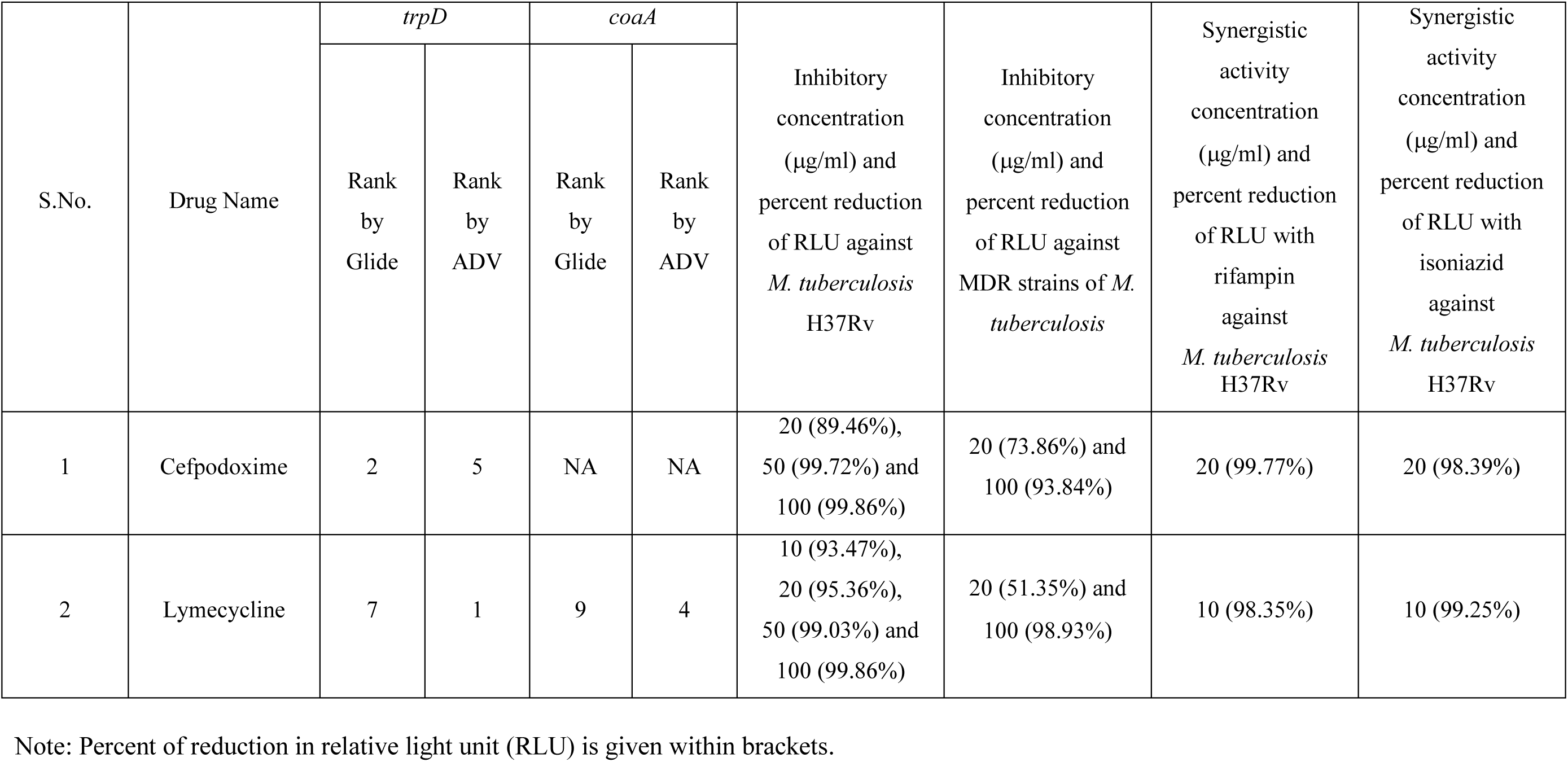
Inhibitory activity of cefpodoxime and lymecycline using luciferase reporter phage assay against *M. tuberculosis*

| S.No. | Drug Name | *trpD* |  | *coaA* |  | Inhibitory concentration (µg/ml) and percent reduction of RLU against *M. tuberculosis* H37Rv | Inhibitory concentration (µg/ml) and percent reduction of RLU against MDR strains of *M. tuberculosis* | Synergistic activity concentration (µg/ml) and percent reduction of RLU with rifampin against *M. tuberculosis* H37Rv | Synergistic activity concentration (µg/ml) and percent reduction of RLU with isoniazid against *M. tuberculosis* H37Rv |
| --- | --- | --- | --- | --- | --- | --- | --- | --- | --- |
|  |  | Rank by Glide | Rank by ADV | Rank by Glide | Rank by ADV |  |  |  |  |
| 1 | Cefpodoxime | 2 | 5 | NA | NA | 20 (89.46%),<br>50 (99.72%) and<br>100 (99.86%) | 20 (73.86%) and<br>100 (93.84%) | 20 (99.77%) | 20 (98.39%) |
| 2 | Lymecycline | 7 | 1 | 9 | 4 | 10 (93.47%),<br>20 (95.36%),<br>50 (99.03%) and<br>100 (99.86%) | 20 (51.35%) and<br>100 (98.93%) | 10 (98.35%) | 10 (99.25%) |

Note: Percent of reduction in relative light unit (RLU) is given within brackets.

### 3.4 Lymecycline and cefpodoxime show synergistic activity with rifampin and isoniazid against *M. tuberculosis*

Synergistic role of cefpodoxime and lymecycline against *M. tuberculosis* were determined in the presence of rifampicin and isoniazid (Table 3). Cefpodoxime showed synergistic inhibitory role at 20 μg/ml concentration with rifampin (2 μg/ml) and isoniazid (0.2 μg/ml) against *M. tuberculosis.* Cefpodoxime (20 μg/ml) showed 89.46% inhibition of *M. tuberculosis* when used alone while 99.77% and 98.39% when used with rifampin and isoniazid respectively. Similarly, lymecycline alone, in the concentration of 10 μg/ml, inhibited 93.47% of *M. tuberculosis* while 98.35% and 99.25% and in the presence of rifampin and isoniazid respectively. These results clearly indicate that cefpodoxime and lymecycline have synergistic role with rifampin and isoniazid against *M. tuberculosis.* Molecular interactions of lymecycline with *trpD*, cefpodoxime with *trpD*, and lymecycline with *coaA* are displayed in Figures 2, 3 and 5 respectively.

**Figure 2:**
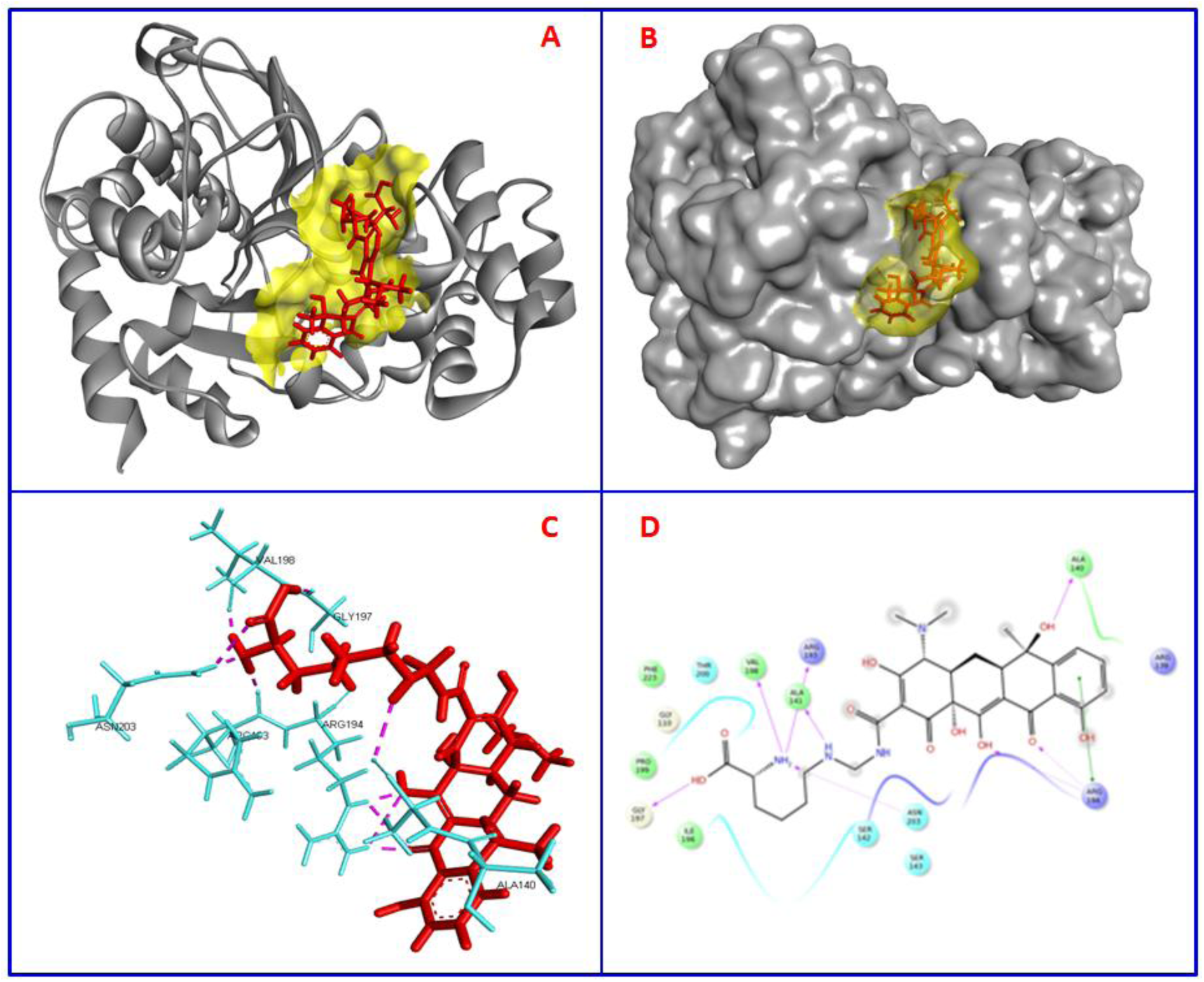
Effective binding of lymecyline in the active site of the *trpD* of *M. tuberculosis*. Fig 2A shows the *trpD* in solid ribbon form (grey in color) while lymecycline is showed in red stick form. The active site of the protein is shown as surface (yellow). Fig 2B shows the overall surface of the *trpD* and lymecycline with the grey color for *trpD* and transparent yellow for lymecycline (stick form). Fig 2C shows the interacting amino acids of *trpD* in the cyan color while the drug lymecycline is displayed in red color stick form with hydrogen bonds in pink color dotted lines. Fig 2D displays the interactions between the *trpD* and lymecycline in two dimensional view.

**Figure 3:**
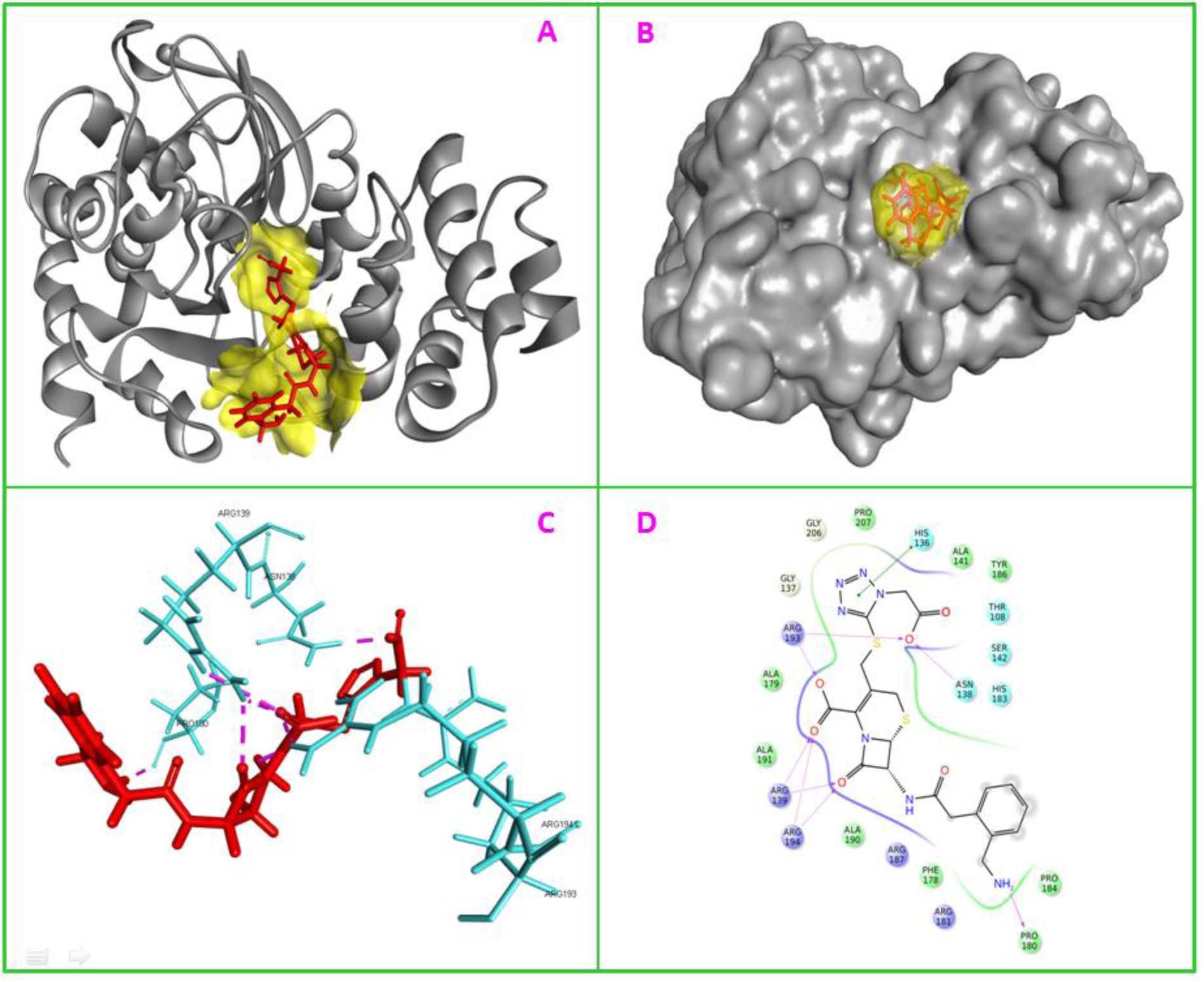
Binding and molecular interactions of cefpodoxime in the active site of the *trpD* of *M. tuberculosis*. Fig 3A shows the *trpD* in solid ribbon form (grey in color) while cefpodoxime is showed in red stick form with the active site of the protein is highlighted as yellow surface. Fig 3B shows the effective fitting of the cefpodoxime in the active site of the *trpD* (yellow surface) and cefpodoxime in stick form. Fig 3C shows the interacting amino acids of *trpD* (cyan color) by hydrogen bonds (pink color dotted lines) with the cefpodoxime (red color, stick form). Fig 3D displays the inter-molecular interactions between the *trpD* and cefpodoxime in two dimensional view.

**Figure4:**
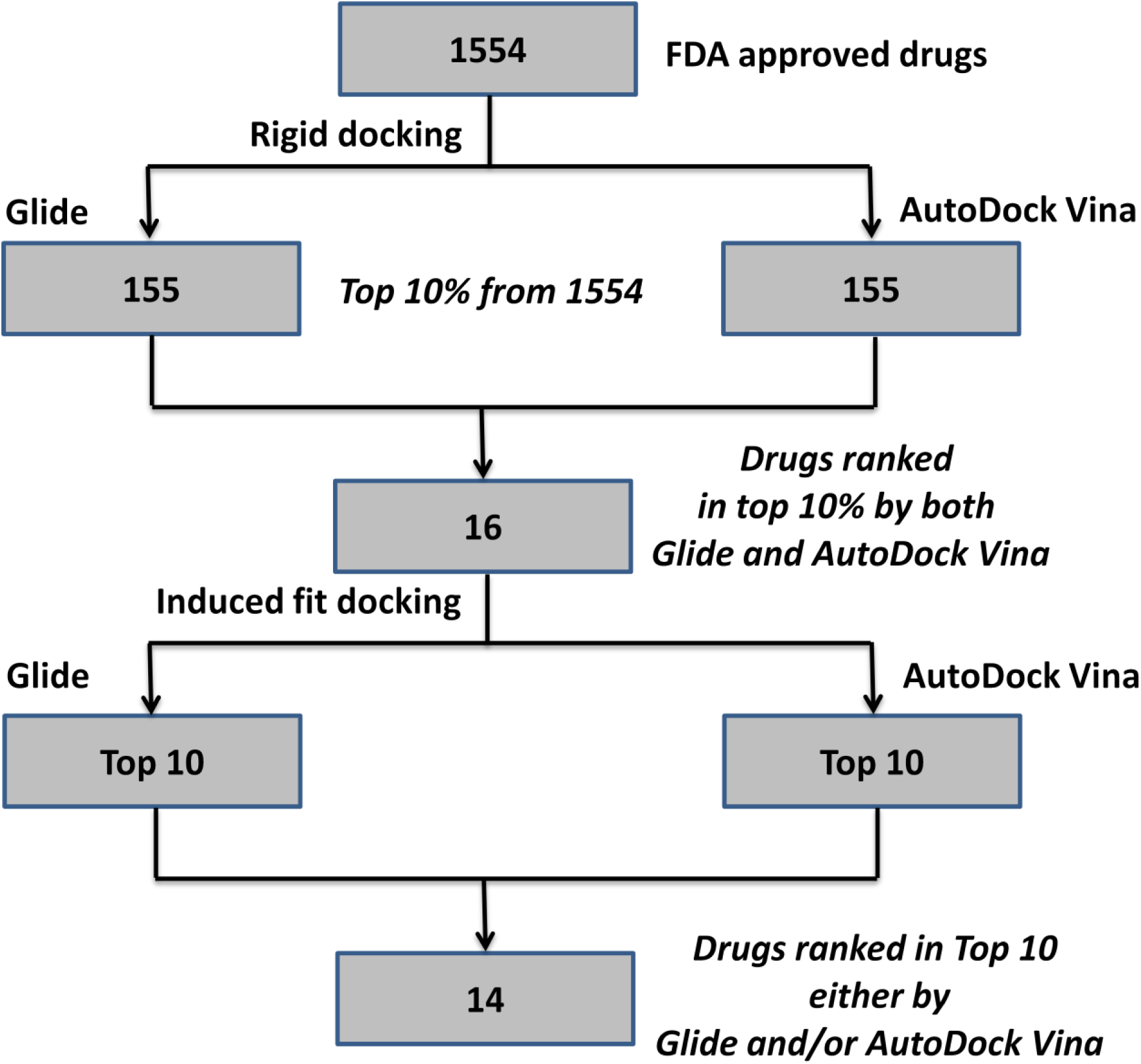
Prioritization of 1554 known drugs using virtual screening against *coaA* of *M. tuberculosis*.

**Figure 5:**
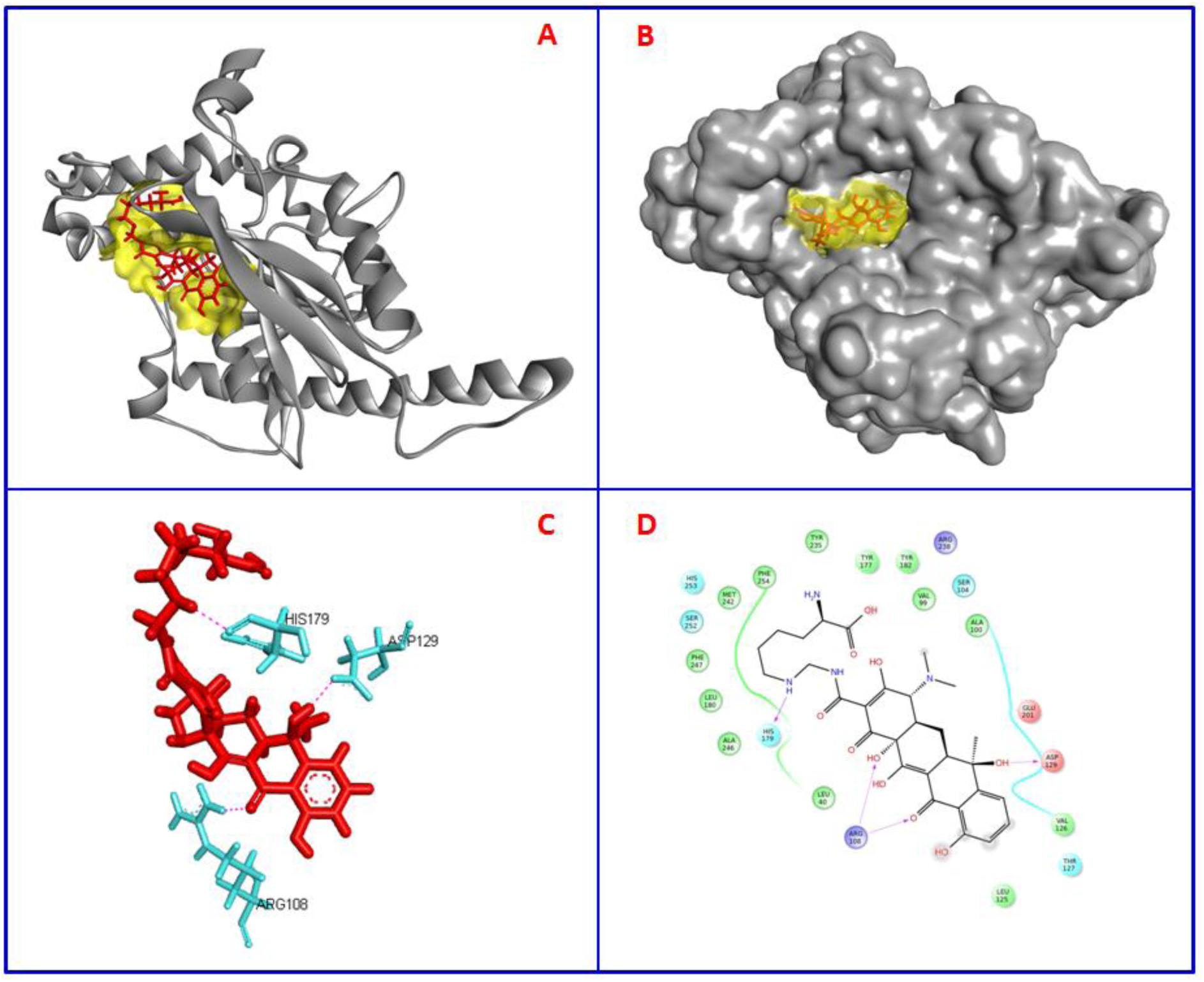
Binding and molecular interactions of lymecycline in the active site of the *coaA* of *M. tuberculosis.* Fig 5A shows the *coaA* in solid ribbon form (grey in color) while lymecycline is showed in red stick form with the active site of the protein is highlighted as yellow surface. Fig 5B shows the effective fitting of the lymecycline in the active site of the *coaA* (yellow surface). Fig 5C shows the interacting amino acids of *coaA* (cyan color) by hydrogen bonds (pink color dotted lines) with the lymecycline (red color, stick form). Fig 5D displays the inter-molecular interactions between the *coaA* and cefpodoxime in two dimensional view.

## 4 DISCUSSION

The long duration of time needed for the treatment of TB using the currently available drugs and the emergence drug resistant TB demand for better therapeutic drugs for the successful and sustained control of TB. Given the cost and time required for the development of new drugs using conventional discovery process, recent success in the repurposing of drugs has encouraged researchers to pursue research in this discipline, especially for TB. Since *M. tuberculosis* encodes for more than 4000 proteins, utmost care should be taken for the selection of target proteins based on which the whole prioritization of drugs is determined. Given the fact that both of the targets selected (*trpD* and *coaA*) are essential for the optimal growth or survival of *M. tuberculosis* (Sassetti et al., 2003; Smith et al., 2001; Awasthy et al., 2010; Sassetti and Rubin, 2003), molecules used to inhibit these proteins are likely to have therapeutic value for tuberculosis.

The *in vitro* activity of cefpodoxime observed in this study is further encouraged by the significant uses of other beta-lactam antibiotics, amoxicillin, imipenem and meropenem in the treatment of DR-TB (WHO, 2014). Cefpodoxime is a semi-synthetic cephalosporin and a beta-lactam bactericidal antibiotic which targets penicillin-binding proteins (PBPs) located in the bacterial cytoplasmic membrane of the target bacteria. Beta-lactum antibiotics are used in combination with clavulanate to get protected from the beta-lactamase of *M. tuberculosis* (Ref: Hugonnet et al., 2009). Out observation of cefpodoxime being active against drug sensitive and resistant *M. tuberculosis* strains indicate its potential role in the treatment of TB. Furthermore, synergistic activity of cefpodoxime with rifampicin as well as with isoniazid suggest that cefpodoxime may potentially reduce the time needed for the treatment of TB in combination with a beta-lactamase inhibitor.

Lymecycline is a tetracycline broad-spectrum antibiotic used for the treatment of urinary tract infection, and other bacterial infections such as gonorrhea and chlamydia. Lymecycline known to bind to the 30S ribosomal subunit and inhibit protein synthesis (https://www.drugbank.ca/). However, the observation of strong binding of lymecycline with *trpD* and coaA and subsequent inhibition of drug sensitive and resistant strains of *M. tuberculosis* in the present study indicates a possible polypharmacology nature of the lymeycline. Further, since lymecycline was found to have synergistic activity with rifampin and isoniazid it may potentially reduce the time needed for the treatment of TB. However, studies in animal models are needed to establish its efficacy against *M. tuberculosis* under *in vivo* conditions.

Our prioritized drugs becomes significant because a total of 4 of these 29 drugs have already been reported to have activity against *M. tuberculosis*. Among the 15 drugs prioritized against *trpD*, two drugs namely acarbose (ranked 1 and 2 by Glide and ADV, respectively) and idarubicin (rank: 10 and 7) were reported previously as active against *M. tuberculosis*. For example, acarbose has been reported to inhibit trehalose synthase (*treS*) of *Mycobacterium smegmatis* which share 83% sequence identity with *treS* of *M. tuberculosis* (Caner et al., 2013; and Pan et al., 2008). Idarubicin is already known to target bacterial primase *dnaG*, a DNA-dependent RNA polymerase and inhibit *M. smegmatis* (MIC = 0.6 μM) and *M. tuberculosis* (MIC = 80 μM) (Gajadeera et al 2015). Similarly, two of the 14 prioritized drugs against *coaA*, namely paromomycin (rank:7 and 5) and daunorubicin (rank: 2 and 14) were previously reported to have antimycobacterial activity. Particularly, paromomycin was demonstrated to have early bactericidal activity against *M. tuberculosis* (Donald et al., 2000) in a pilot clinical research study in newly diagnosed pulmonary tuberculosis patients (Donald et al., 2000). Daunorubicin was recently demonstrated to inhibit growth of *M. tuberulosis* and *M. smegmatis* under *in vitro* conditions (Gajadeera et al., 2015). Since four of 29 prioritized drugs (acarbose, idarubicin, paromomycin and daunorubicin) have already reported to have potential use for the treatment of TB, especially evidence generated from clinical research for paromomycin (Donald et al., 2000) encourage us to speculate that other drugs prioritized using the same method are likely to have potential activity against *M. tuberculosis*.

## Conclusion

Since 4 out of 25 prioritized drugs are already reported to have potential use against *M. tuberculosis* using *in vitro* studies and clinical research, the procedure employed in this study becomes significant, thus, it can be adopted to prioritize drugs for repurposing against any other diseases. Biological activity of lymecycline and cefpodoxime against drug sensitive and drug resistant strains of *M. tuberculosis* and their synergistic role with rifampin and isoniazid suggest that these two drugs have potential for repurposed in the treatment of TB.

## ACKNOWLEDGEMENTS

SB acknowledges sincerely the management of Loyola College, Chennai, India for providing the facility to carry out her Ph.D work. JCS is supported by ICMR-Biomedical Informatics Centre, NIRT, Chennai, India.

